# Hydration Energetics Shape Antibody Discrimination between Sulfotyrosine and Phosphotyrosine

**DOI:** 10.64898/2026.08.05.743142

**Authors:** Takahiro Mori, Kotaro Yahagi, Saki Maruoka, Kota Toyoda, Yoshikazu Sonoshita, Yohei Kametani, Yoshihito Shiota, Kazunari Yoshizawa, Keiichi Watanabe, Kyo Okazaki, Yoshihiro Kobashigawa, Hiroshi Morioka, Hideki Hirakawa, Etsuko Nishimoto, Takamasa Teramoto, Yoshimitsu Kakuta

## Abstract

Chemically similar post-translational modifications can mediate distinct biological functions, but how proteins distinguish between them remains unclear. Sulfotyrosine (sTyr) and phosphotyrosine (pTyr) exemplify this problem because they have similar sizes, local geometries, and electrostatic properties but function in different biological contexts. Here, we used the monoclonal antibody PSG2, which recognizes sTyr independently of the surrounding peptide sequence, to examine how a protein distinguishes these modifications. The crystal structure of PSG2 bound to an sTyr-containing peptide revealed a deep electropositive pocket with no modeled water molecules in direct contact with the sulfate group. Gas-phase density functional theory calculations favored pTyr over sTyr, showing that direct protein–ligand interactions alone are insufficient to explain PSG2 selectivity. Explicit first-shell hydration calculations showed that pTyr has a larger desolvation penalty than sTyr, and accounting for this difference reversed the calculated energetic order. Isothermal titration calorimetry showed favorable enthalpic and entropic contributions to sTyr binding, whereas no detectable heat signal was observed for pTyr. These results show that PSG2 distinguishes sTyr from pTyr through the balance between direct protein–ligand interactions and ligand desolvation.

## Introduction

The selective recognition of biomolecules underlies essential processes such as enzyme catalysis, immune surveillance, and cellular signaling^1,2^. At the molecular level, specificity is generally explained by steric and electrostatic complementarity^3^, principles firmly established by structural biology. Classic examples include enzymes that distinguish between distinct substrates and antibodies that bind antigens via shape and charge complementarity^4,5^.

While these cases are well understood, many biologically crucial events require discrimination between molecules that are chemically very similar. Ion channels, for example, can distinguish between K^+^ and Na^+^ ions by exploiting subtle differences in desolvation energetics^6–8^. Carbohydrate-binding proteins can recognize stereoisomers that differ only by the orientation of a single hydroxyl group; RNA-binding proteins detect nucleotide modifications within otherwise identical sequences; and metalloenzymes achieve selectivity among ions of comparable size and charge^9–11^.

Unlike these relatively well-characterized systems, distinguishing sulfate (-SO_3_^−^) from phosphate (-PO_3_^2−^/-HPO_3_^−^) groups remains difficult. Despite their similar tetrahedral geometry and related electrostatic features, sulfate and phosphate confer distinct biological functions when covalently attached to proteins, glycans, or metabolites^12–16^. At the level of post-translational modifications, sulfotyrosine (sTyr) and phosphotyrosine (pTyr) epitomize the challenge of distinguishing sulfate from phosphate groups^17^. Both modify the phenolic hydroxyl group of tyrosine and share closely related local geometries and electrostatic features, while differing in protonation and solvation properties^16,18^ (Figure 1a and Figure S1). However, they function in different biological contexts. Sulfation, catalyzed by tyrosylprotein sulfotransferases (TPSTs) utilizing 3′-phosphoadenosine-5′-phosphosulfate (PAPS), typically modifies extracellular and membrane-associated proteins^19–21^. This modification forms stable recognition motifs that mediate processes such as chemokine signaling, leukocyte adhesion, viral entry, and sulfation-dependent anticoagulant activity^14,22–24^. Structural studies of human TPST–substrate complexes and the tick TPST–madanin complex have revealed how TPST enzymes recognize tyrosine-containing substrates and catalyze site-specific tyrosine sulfation, providing a molecular basis for the generation of these functional sulfotyrosine motifs^25^. For instance, sulfotyrosine residues on the chemokine receptor CCR5 are essential for HIV-1 entry into host cells^24,26^. In contrast, phosphorylation is an intracellular modification catalyzed by protein tyrosine kinases (PTKs)^27^. It is reversible and functions as a dynamic molecular switch in signaling cascades controlling proliferation, differentiation, and metabolism by recruiting specific phosphotyrosine-binding domains, such as SH2 domains^28,29^.

**Figure 1.**
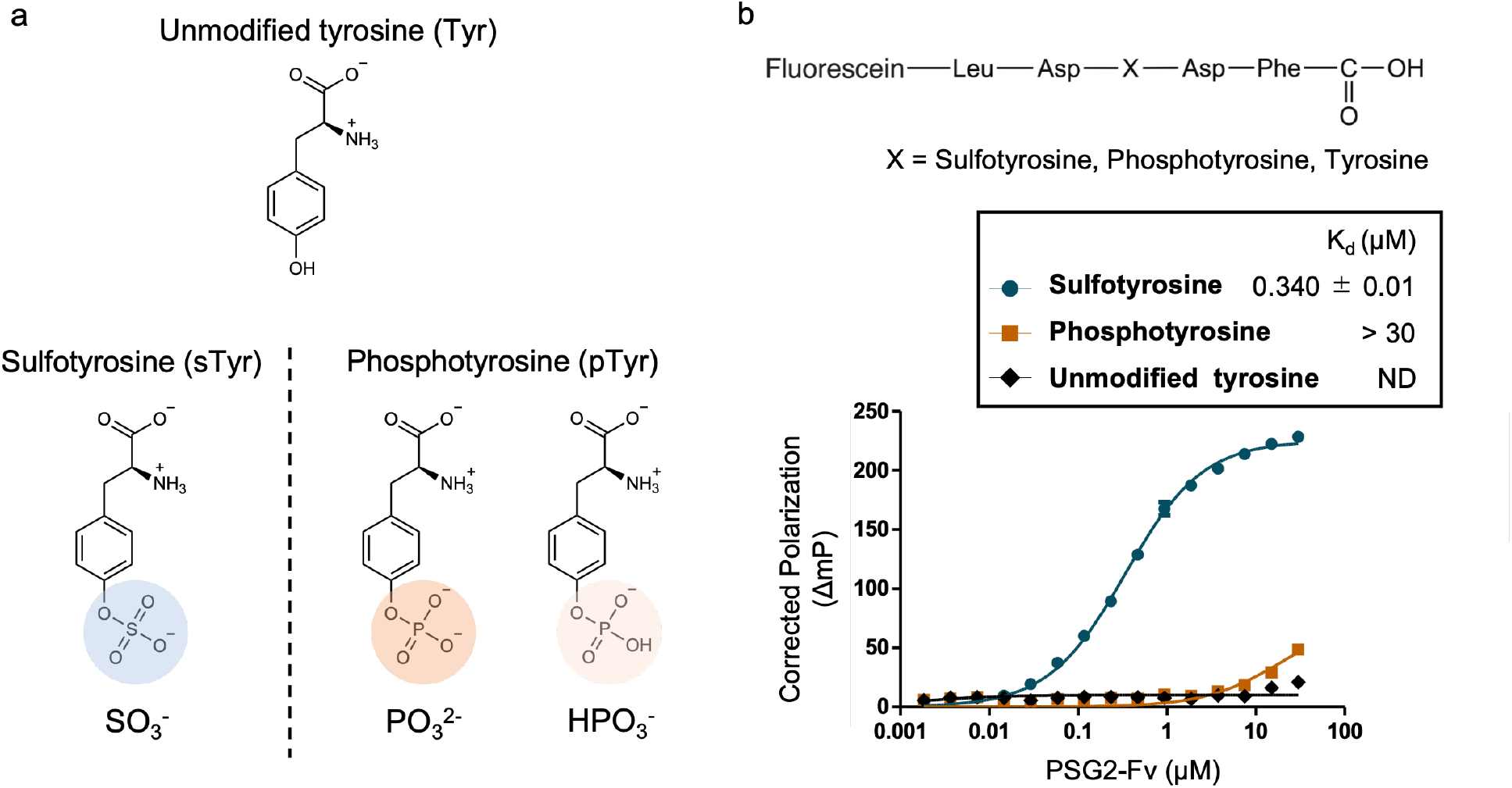
(a) Structural representations of unmodified tyrosine, sulfotyrosine (SO_3_^−^), dianionic phosphotyrosine (PO_3_^2−^), and monoanionic phosphotyrosine (HPO_3_^−^). (b) Top: Fluorescently labeled peptide used in binding assays, incorporating sulfotyrosine (sTyr), phosphotyrosine (pTyr), or unmodified tyrosine (Tyr) at the central position. Bottom: Fluorescence polarization measurements of PSG2-Fv binding to a sulfotyrosine-containing peptide, a phosphotyrosine-containing peptide, and an unmodified tyrosine-containing peptide. Data are shown as means ± SEM from triplicate measurements.

These biological differences have motivated the development of methods for detecting tyrosine sulfation and distinguishing it from phosphorylation. Recent studies have developed chemical methods for controlling tyrosine sulfation, as well as engineered binding domains, monoclonal antibodies, and nanopore-based sensors for detecting sulfotyrosine^30–34^. Although these approaches have improved the detection of tyrosine sulfation, few studies have quantitatively examined the physicochemical factors that allow sTyr to be distinguished from pTyr at the atomic level. The difficulty of distinguishing these modifications is illustrated by the ongoing debate over histone tyrosine sulfation. Recent studies have proposed regulatory roles for this modification^35^. However, these findings remain controversial because pTyr may have been misidentified as sTyr in mass spectrometry and antibody-based assays^36,37^. This debate shows that the close physicochemical similarity between sTyr and pTyr can complicate their experimental identification and lead to different biological interpretations. Underlying these challenges is the limited understanding of how biological macromolecules, including antibodies, enzymes, and receptor domains, distinguish sulfate from phosphate groups.

PSG2 is a monoclonal antibody that recognizes sTyr independently of its surrounding sequence^31^. Here, we used X-ray crystallography, density functional theory (DFT), and isothermal titration calorimetry (ITC) to investigate the molecular basis of its selectivity. Our results show that shape and charge complementarity alone cannot explain the discrimination between sTyr and pTyr. Instead, differences in the desolvation penalties of sulfate and phosphate contribute to this selectivity. These findings provide a physicochemical basis for the discrimination between sTyr and pTyr by PSG2.

## Results

### PSG2 specifically recognizes sulfotyrosine

To enable detailed structural and functional characterization, we engineered a stabilized Fv-clasp form of the PSG2 antibody (PSG2-Fv), facilitating efficient recombinant expression in *Escherichia coli*^38^ (Figure S2 and Table S1). This Fv-clasp construct includes a coiled-coil domain designed to ensure correct pairing and folding of the heavy and light chains, thereby preserving antigen-binding functionality. The observed melting temperature (T_m_) of PSG2-Fv in the absence of ligand was 49.7 °C (Figure S3), which is within the expected range for properly folded antibody Fv-clasp constructs expressed in *E. coli*^38^, suggesting that the Fv-clasp adopts a stable, native-like conformation. We evaluated the specificity of PSG2-Fv using fluorescence-based binding assays with peptides containing sulfotyrosine, phosphotyrosine, or unmodified tyrosine residues previously described by Hoffhines *et al*.^31^ (Figure 1b and Table S2). PSG2-Fv bound the sTyr-containing peptide with high affinity (K_d_ = 0.340 ± 0.01 μM), whereas pTyr- and unmodified Tyr-containing peptides showed no saturable binding under the conditions tested.

### Structural basis of sulfotyrosine recognition by PSG2

To examine how PSG2 recognizes sTyr, we determined the crystal structure of PSG2-Fv bound to an untagged sulfotyrosine-containing peptide (Leu-Asp-Tyr(-SO_3_^−^)-Asp-Phe) at 2.48 Å resolution (Figure 2 and Table S3). The electron-density maps allowed the sTyr residue to be modeled, whereas the density for the adjacent peptide residues was less well defined, particularly for the Leu residue (Figure 2b).

**Figure 2.**
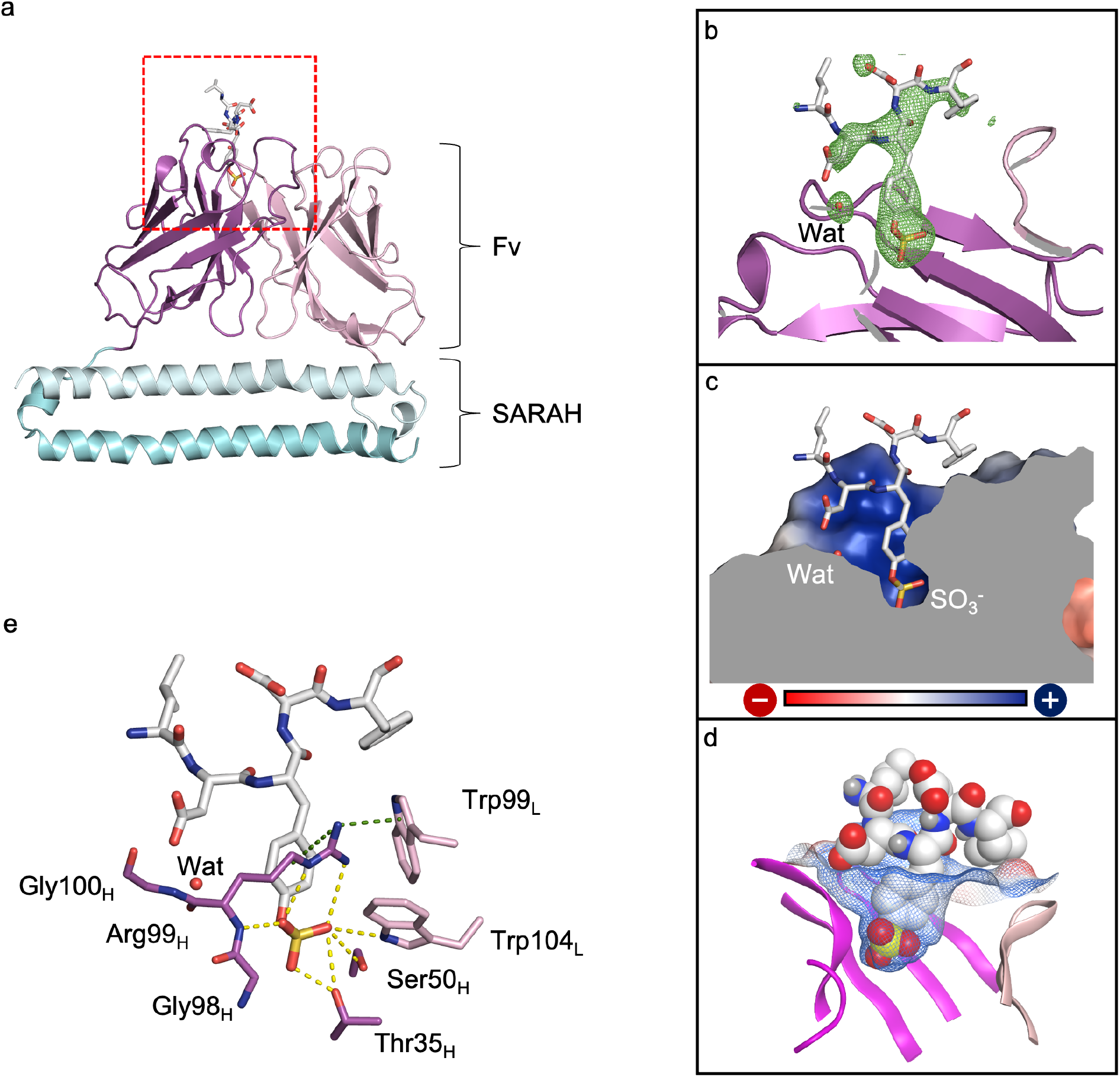
(a) Overall ribbon model of the PSG2-Fv in complex with a sulfotyrosine-containing peptide. The light chain is colored light pink, the heavy chain is magenta, and the coiled-coil SARAH domain is light blue. The sulfotyrosine-containing peptide is shown in white sticks. (b) Polder map of the sulfotyrosine-containing peptide (contour level = 2.5σ) and water molecule (contour level = 3σ). (c) Cutaway view of the PSG2 binding pocket, with the electrostatic surface potential mapped onto the molecular surface (blue: positive; red: negative). The bound sulfotyrosine is shown in stick model. (d) Molecular surface representation of PSG2-Fv. The molecular surface is depicted as a mesh, while the sulfotyrosine-containing peptide is represented as spheres based on Van der Waals radii. (e) Close-up view of the antigen recognition site. The sulfotyrosine-containing peptide is shown in stick model. The light chain of PSG2-Fv is colored in light pink and the heavy chain of PSG2-Fv is colored in magenta. Cation-π interactions are indicated by green dashed lines, and hydrogen bonds are shown by yellow dashed lines.

The antigen-binding site of PSG2-Fv comprises a deep, electropositive binding pocket specifically suited to accommodate the negatively charged sulfate group of sulfotyrosine (Figure 2c). The sulfate moiety participates in multiple hydrogen-bonding interactions with side chains of heavychain residues Thr35 and Ser50, the main-chain nitrogen and side chain of heavy chain Arg99, and the side chain of light chain Trp104 (Figure 2e). The aromatic ring of sTyr is also positioned near the guanidinium group of Arg99, forming a cation–π interaction (Figure S4).

No water molecules were modeled in direct contact with the sulfate group, suggesting that little hydration water is retained around the sulfate group in the bound state. A single water molecule was resolved, bridging the peptide’s Asp residue and the backbone of Arg99 (Figure 2b and 2d). Based on the structure of the sTyr-bound complex, an un-modified tyrosine residue would be expected to provide less favorable shape and charge complementarity with the PSG2 binding pocket. This interpretation is consistent with the absence of saturable binding to the unmodified tyro-sine-containing peptide under the conditions tested. Together, the deep electropositive pocket and the hydrogen-bonding and cation–π interactions explain the structural recognition of sTyr by PSG2-Fv.

### Gas-phase binding energies do not explain the sulfotyrosine selectivity of PSG2

The crystal structure explained why unmodified Tyr is unlikely to form the same interactions as sTyr but did not explain how PSG2-Fv distinguishes sTyr from pTyr. Removal of the sulfate group eliminates the hydrogen-bonding interactions observed in the sTyr-bound structure. In contrast, phosphate has a tetrahedral geometry and charge distribution similar to those of sulfate and may therefore form comparable hydrogen-bonding interactions within the electropositive binding pocket. We therefore examined whether the PSG2-Fv binding site energetically favors pTyr over sTyr in the absence of solvent effects (Figure 3a).

**Figure 3.**
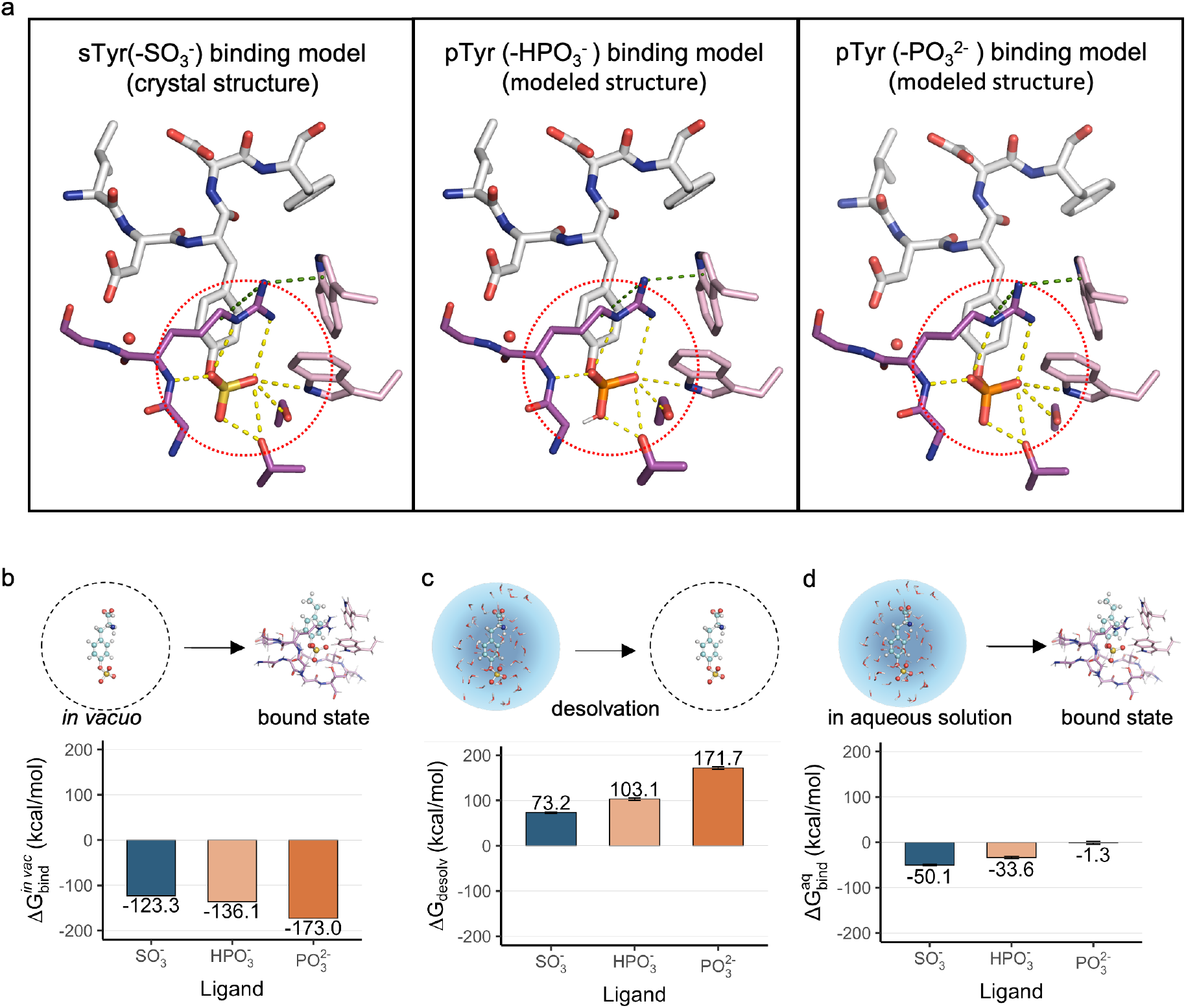
(a) Close-up views of the PSG2-Fv binding pocket showing sulfotyrosine (left, crystal structure), monoanionic phosphotyrosine (center, modeled structure), and dianionic phosphotyrosine (right, modeled structure). Atoms are colored as follows: carbon and hydrogen (white), oxygen (red), nitrogen (blue), sulfur (yellow), and phosphorus (orange). Green dashed lines indicate cation– π interactions; yellow dashed lines indicate hydrogen bonds. (b) Vacuum-phase binding free energies (kcal/mol) for PSG2-Fv complexes with sulfotyrosine (SO_3_^−^) and phosphotyrosine (PO_3_^2−^ and HPO_3_^−^), showing stronger predicted interactions for pTyr than sTyr, contrary to experimental findings. (c) Estimated dehydration penalties (kcal/mol) for each ligand. Bars show means; error bars denote 95% confidence intervals. (d) Hydration-corrected cluster-model energies calculated using ligand·(H^2^O)_11_ clusters constructed from the 11 nearest water molecules in MD simulations.

We first performed gas-phase quantum chemical calculations based on the crystal structure of the PSG2-Fv–sTyr complex, without explicitly modeling water molecules. We created pTyr binding models by replacing the sulfate group with a phosphate group. Given that pTyr has a pKa of 6.1^39^, it exists predominantly in the dianionic form (-PO_3_^2−^) but also partially in the monoanionic form (-HPO_3_^−^) at physiological cytosolic pH (∼7.2)^40^. For monoanionic pTyr, all three possible protonation-site isomers were independently optimized within the PSG2-Fv binding-site model, and the isomer with the lowest Gibbs free energy was used for subsequent calculations. We then calculated the dissociation-limit binding energies of the crystal-structure-based sTyr model and the two modeled pTyr complexes, together with their corresponding isolated ligands (Figure S5). The calculated gas-phase binding energies were -123.3 kcal mol^−1^ for the sTyr model, -136.1 kcal mol^−1^ for the Tyr– HPO_3_^−^ model, and -173.0 kcal mol^−1^ for the Tyr–PO_3_^2−^ model (Figure 3b). Thus, both pTyr models showed more favorable gas-phase binding energies than the sTyr model, contrary to the fluorescence polarization results. These results indicate that gas-phase binding energies alone are insufficient to explain the observed selectivity of PSG2 for sTyr.

### Ligand desolvation contributes to the sTyr selectivity of PSG2

The disagreement between the fluorescence polarization results and the gas-phase quantum chemical calculations prompted us to examine the effects of solvation, which were not included in the *in vacuo* models. Because the crystal structure was determined at 2.48 Å resolution, not all water molecules could be identified with confidence. We therefore calculated the water-density distribution around the sulfate group using the three-dimensional reference interaction site model (3D-RISM).^41^ Applying a density threshold of 4.0 relative to bulk water, the analysis indicated no stable water sites within 4 Å of the sulfotyrosine (Figure S6). Together with the absence of modeled water molecules in direct contact with the sulfate group in the crystal structure, this result suggests that little hydration water is retained around the sulfate group in the bound state (Figure 2d and e). Previous studies have shown that sulfate and phosphate ions differ in their hydration properties^42,43^. We therefore examined whether the corresponding groups in sTyr and pTyr also differ in their desolvation penalties.

To test this hypothesis, we constructed explicit first-shell hydration models for sTyr, Tyr–HPO_3_^−^, and Tyr–PO_3_^2−^. For each ligand, the 11 water molecules closest to the sulfate or phosphate group were extracted from equilibrated molecular dynamics trajectories (Figure S7). This number was selected to represent a saturated first hydration shell, based on coordination numbers of approximately 11 reported for sulfate and phosphate ions in *ab initio* molecular dynamics studies^43,44^. Quantum chemical calculations were then performed for each hydrated ligand–water cluster, the corresponding isolated ligand, and the associated water cluster. The energy differences between these states were used to evaluate the stabilization provided by the first hydration shell and the desolvation penalty associated with loss of the ligand–water interactions. The calculated desolvation penalties were 73.2 kcal mol^−1^ for sTyr, 103.1 kcal mol^−1^ for Tyr–HPO_3_^−^, and 171.7 kcal mol^−1^ for Tyr–PO_3_^2−^ (Figure 3c and Figure S8), increasing in the order sTyr < Tyr–HPO_3_^−^ < Tyr–PO_3_^2−^. We next combined the calculated desolvation penalties with the corresponding gas-phase binding energies. Although the gas-phase calculations favored Tyr– HPO_3_^−^ and Tyr–PO_3_^2−^ over sTyr, accounting for ligand desolvation reversed the energetic order. The resulting hydration-corrected energies were -50.1 kcal mol^−1^ for sTyr, - 33.6 kcal mol^−1^ for Tyr–HPO_3_^−^, and -1.3 kcal mol^−1^ for Tyr– PO_3_^2−^ (Figure 3d). Thus, the larger desolvation penalties of the two pTyr forms offset their more favorable gas-phase interactions and resulted in an energetic preference for sTyr. These results identify differences in ligand desolvation penalties as a factor contributing to the sTyr selectivity of PSG2.

Finally, we used isothermal titration calorimetry (ITC) to characterize the thermodynamics of PSG2-Fv binding to an untagged sTyr-containing peptide, Asp-Tyr(−SO_3_^−^)-Asp (Figure 4 and Table 1). Binding was favorable in both enthalpic and entropic terms, with ΔH = −5.5 kcal mol^−1^ and −TΔS = −2.1 kcal mol^−1^. The favorable enthalpic contribution is compatible with the direct interactions observed in the crystal structure, whereas the favorable entropic contribution is compatible with solvent release upon binding, although other contributions to the entropy change cannot be excluded. In contrast, titration with the corresponding pTyr-containing peptide, Asp-Tyr(−PO_3_^2−^)-Asp, produced no detectable heat signal under the measurement conditions. Thus, favorable binding thermodynamics were detected for sTyr but not for pTyr.

**Table 1.** Thermodynamic parameters of the association of a sulfotyrosine-containing peptide with PSG2-Fv. Values are means; 95% confidence intervals are shown in brackets. Parameters refer to the first binding site; n is the binding stoichiometry.

| *n* | *K<sub>d</sub>* (μM) | Δ*G* (kcal/mol) | Δ*H* (kcal/mol) | -*T*Δ*S* (kcal/mol) |
| --- | --- | --- | --- | --- |
| 1.13 [0.92; 1.34] | 2.79 [0.40; 5.19] | -7.60 [-8.17; -7.03] | -5.52 [-5.81; -5.24] | -2.09 [-2.89; -1.30] |

**Figure 4.**
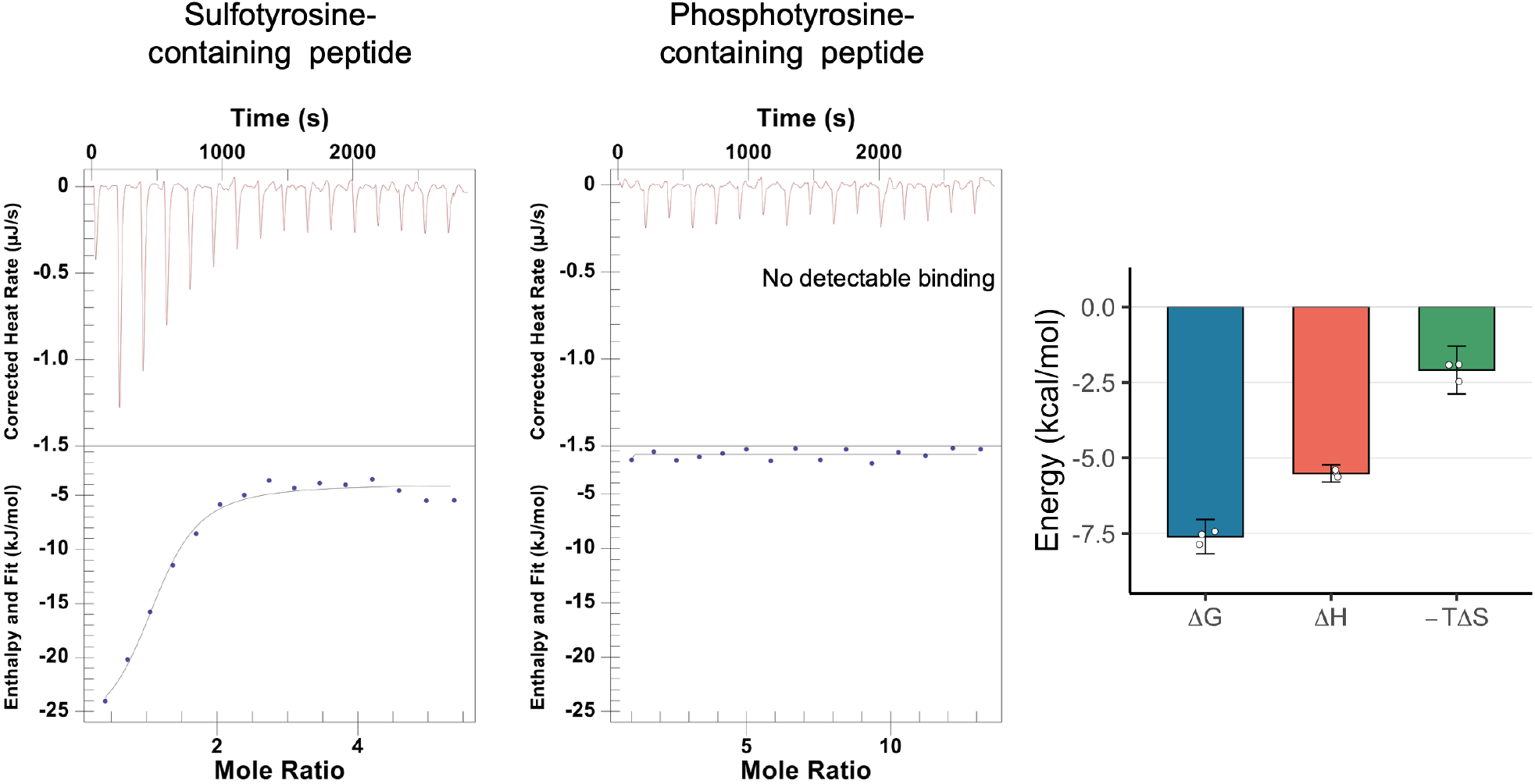
ITC analysis of the interaction between PSG2-Fv and a sulfotyrosine-containing peptide (Asp-Tyr(SO_3_^−^)-Asp) (left) or a phosphotyrosine-containing peptide (Asp-Tyr(PO_3_^2−^)-Asp) (right). The bar graph summarizes thermodynamic parameters for sTyr binding only. Bars show means; error bars denote 95% confidence intervals across independent ITC replicates; points are individual experiments.

## Discussion

The ITC data showed that PSG2-Fv binding to sTyr is favored by both enthalpy and entropy. The favorable enthalpic contribution can be explained in part by the hydrogen bonds and cation–π interaction observed in the crystal structure. Although the entropy change cannot be assigned solely to solvent release, the favorable entropic contribution may partly reflect the release of water from the ligand and binding pocket. This interpretation agrees with the water-poor binding environment indicated by the crystal structure and 3D-RISM analysis. In contrast, no detectable heat signal was observed for pTyr under the same conditions.

The selectivity of PSG2 cannot be explained by the interactions formed in the binding pocket alone. The gas-phase calculations favored both forms of pTyr over sTyr, showing that the phosphate group can form favorable interactions with the PSG2 binding site. However, pTyr is more strongly stabilized by its hydration shell and has a larger desolvation penalty than sTyr. To bind to the water-poor PSG2 pocket, pTyr would need to lose part of these favorable hydration interactions. The energy required for this loss offsets the favorable interactions that pTyr can form with the protein. In contrast, sTyr can bind to the same pocket with a smaller desolvation penalty. PSG2 therefore favors sTyr through the balance between direct protein–ligand interactions and ligand desolvation (Figure S9).

Differences in desolvation penalties also contribute to the discrimination of chemically similar ions. Potassium channels such as KcsA and MthK distinguish K^+^ from Na^+^ partly because the two ions require different amounts of energy to lose their hydration water^6,8^. Our results show that a similar mechanism can apply to covalently attached post-translational modifications. In both cases, selectivity depends not only on the interactions formed after binding but also on the energy required to remove hydration water. It remains unclear whether other sTyr-binding proteins also use differences in desolvation penalties to distinguish sTyr from pTyr.

Comparison with the anti-pTyr antibody 4G10 shows how the hydration environment of a binding pocket may affect ligand preference. The 4G10 structure contains a shallow, solvent-exposed pocket in which the phosphate group interacts with several basic residues while retaining water molecules (Figure S10)^45^. This binding mode may allow pTyr to retain part of its hydration shell and reduce its desolvation penalty. PSG2 instead contains a deeper, water-poor pocket with no modeled water molecules in direct contact with the sulfate group. In this environment, desolvation of the more strongly hydrated phosphate group requires more energy than desolvation of the sulfate group. The different hydration environments of these two antibodies may therefore contribute to their opposite ligand preferences.

These findings explain how PSG2 distinguishes between the chemically similar sulfate and phosphate groups of sTyr and pTyr. Shape and charge complementarity in the bound state alone are not sufficient. The water-poor PSG2 pocket favors sTyr because its sulfate group has a smaller desolvation penalty than the phosphate group of pTyr. PSG2 therefore distinguishes these modifications through both direct protein–ligand interactions and the energetic cost of ligand desolvation. This mechanism may also affect the specificity and cross-reactivity of other proteins and analytical reagents that distinguish sulfate from phosphate groups.

## ASSOCIATED CONTENT

### Supporting Information

The Supporting Information is included in this PDF.

## Author Contributions

The manuscript was written through contributions of all authors. All authors have given approval to the final version of the manuscript. Takahiro Mori performed most of the experiments and wrote the initial manuscript. Kotaro Yahagi, Saki Maruoka, Kota Toyoda, and Etsuko Nishimoto performed part of the experiments. Yoshikazu Sonoshita, Yohei Kametani, Yoshihito Shiota, Kazunari Yoshizawa, and Keiichi Watanabe contributed to the DFT calculations. Kyo Okazaki, Yoshihiro Kobashigawa, Hiroshi Morioka, and Hideki Hirakawa contributed to the ITC measurements. Takamasa Teramoto contributed to the experimental work and final manuscript revision. Yoshimitsu Kakuta conceived and supervised the project, directed the overall study, and finalized the manuscript.

## ACKNOWLEDGMENT

The synchrotron radiation experiments were performed at SPring-8 (proposal nos. 2021B2553 and 2022A2553). The computations were carried out using computer resources provided under the General Projects category by the Research Institute for Information Technology, Kyushu University. This work was supported by JST SPRING, Grant Number JPMJSP2136 (to T.M.), and JSPS KAKENHI Grant Numbers JP24K21245 (to K.Y.), JP24K09353 and JP25H01276 (to Y.K.), JP26KJ1809 (to T.M.). During the preparation of this manuscript, ChatGPT (OpenAI) was used solely to assist with English-language editing. The authors reviewed and edited the manuscript and take full responsibility for its content.

## Supporting Information

### Methods

#### Protein expression and purification

For construction of the Fv-clasp expression plasmids, synthetic DNAs encoding SARAH-domain sequences fused downstream of the VH and VL cassettes were prepared based on a previous report^1^. The VH-SARAH and VL-SARAH coding sequences of PSG2-Fv were separately subcloned into the pE_His vector, which adds an N-terminal Ser-Lys-Ile-Lys-His_6_ tag (Table S1). The VH-SARAH and VL-SARAH chains were separately overexpressed in *E. coli* BL21-CodonPlus (DE3)-RIL cells (Agilent Technologies) harboring the corresponding plasmids. The cells were grown to an OD_600_ of 0.4-0.6 and induced with 0.5 mM IPTG overnight at 37°C. The cells were collected by centrifugation, and the resulting pellets were resuspended in a lysis buffer containing 50 mM Tris-HCl and 500 mM NaCl (pH 8.0) and then stored at –80 °C until use. The cells were lysed by sonication and the lysate was collected by centrifugation. The inclusion bodies were isolated by centrifugation and solubilized in a solution of 50 mM Tris, 150 mM NaCl, 6 M guanidine-HCl, and 375 μM β-mercaptoethanol at pH 8.0. Denatured and solubilized VH-SARAH and VL-SARAH chains were mixed in roughly equimolar ratios, and the denaturing reagent was diluted in two steps to promote protein folding. The first step was dilution with 4 M urea, 0.4 M L-arginine, 375 μM GSSG, 50 mM Tris-HCl, and 150 mM NaCl at pH 8.0, and the second step was dilution with 0.4 M L-arginine, 375 μM GSSG, 50 mM Tris-HCl, and 150 mM NaCl at pH 8.0. The protein was purified further using a Superdex 200 16/60 pg column (Cytiva), equilibrated with 50 mM Tris-HCl and 150 mM NaCl at pH 8.0. PSG2-Fv was eluted from the column as a monodisperse peak. Subsequently, it was purified further on a HiTrap Q column equilibrated with 20 mM Tris (pH 7.5) via anion exchange. Bound proteins were then eluted with a linear gradient from 0 M to 1 M NaCl in 20 mM Tris-HCl (pH 7.5). The peak fractions containing target proteins were pooled and concentrated. The final purified PSG2-Fv was concentrated to 9.4 mg/mL.

#### Fluorescence polarization binding assays

Binding assays were performed in LBS OptiPlate-384 F black polystyrene plates (PerkinElmer). PSG2-Fv was serially diluted over a concentration range of 1.5 nM to 25 μM. PSG2-Fv–ligand mixtures were incubated at 23 ± 1 °C in 50 mM Tris-HCl buffer (pH 7.5) containing 200 mM NaCl and 2 nM of an N-terminally fluorescein-labeled peptide containing sulfotyrosine, phosphotyrosine, or unmodified tyrosine. Fluorescence polarization was measured using a Tecan Ultra plate reader with excitation and emission wavelengths of 485 and 535 nm, respectively. Measurements were performed in triplicate at each PSG2-Fv concentration using PSG2-Fv obtained from a single expression and purification batch. Data are presented as means ± SEM. Binding curves were analyzed by nonlinear regression using a one-site binding model in GraphPad Prism 5, and dissociation constants (K_d) were estimated from the fitted curves.

#### Crystallization, data collection, structure determination, and refinement

Crystals of the PSG2-Fv–sulfotyrosine peptide complex were obtained using the hanging drop vapor diffusion method at 20°C. The hanging drops were prepared by combining 1 µL of protein solution (9.4 mg/mL) with 2 mM sulfotyrosine-containing peptide, and 1 µL reservoir solution (3.8 M Sodium formate). The resulting crystals were flash frozen in liquid nitrogen. The X-ray diffraction data were collected using the beamline BL45XU (SPring-8, Hyogo, Japan) system with a wavelength of 1.000 Å. The data were processed using the ZOO system^2^. Molecular replacement for phase determination was performed using Phaser ^3^ with search models such as AlphaFold2 prediction model^4^. The model was constructed using the Coot program ^5^. A refined structure was obtained using the Phenix.refine program ^6^. The resulting structures exhibited good geometries, as assessed using MolProbity ^7^. The atomic coordinates and structure factors of the crystal structure have been deposited in the Protein Data Bank (PDB).

#### 3D-RISM calculations

3D-RISM calculations were performed using Molecular Operating Environment (MOE) Solvent Analysis (Binding mode) ^8,9^. Receptor atoms within 10 Å of the ligand were included. The solvent box used a 10 Å buffer (minimum distance from any solute atom to the box edge) with 0.30 Å grid spacing in each direction. Convergence was set to Medium with NDIIS = 5; Salt and Hydrophobe densities were set to None. Default parameters were used for temperature and ionic strength. Maps were analyzed/visualized in the Solvent-Analysis – Grids panel. To assess the hydration structure, the 3D spatial distribution of water oxygen density around the sulfate group was analyzed. The calculated values represent the local probability of finding a water molecule relative to the bulk solvent density. For visualization, isosurfaces defined by a relative density threshold of 4.0 (four times the bulk density) were rendered within 4 Å of the sulfotyrosine residue to visualize high-density hydration regions.

#### Construction of tyrosine- and phosphotyrosine-containing peptide models

Molecular models of PSG2-Fv bound to tyrosine- or phosphotyrosine-containing peptides were constructed using MOE. The PSG2-Fv–sulfotyrosine complex was used as the template. The sulfotyrosine residue in the bound peptide was replaced with either unmodified tyrosine or dianionic phosphotyrosine, corresponding to −PO_3_^2−^, to generate tyrosine- and phosphotyrosine-containing peptide models, respectively. The resulting models were used to compare possible binding modes and interactions of tyrosine- and phosphotyrosine-containing peptides in the PSG2-Fv binding pocket.

#### Construction of PSG2-Fv–phosphotyrosine binding-site models for DFT calculations

Molecular models of PSG2-Fv bound to phosphotyrosine were constructed using MOE. The phosphotyrosine-bound models were generated from the PSG2-Fv–sulfotyrosine complex by using the sulfotyrosine position as a template for placing phosphotyrosine in the binding pocket. Phosphotyrosine was modeled as an isolated modified tyrosine ligand. Both monoanionic and dianionic phosphotyrosine states, corresponding to −HPO_3_^−^ and −PO_3_^2−^, respectively, were prepared. For the monoanionic state, three protonation-site isomers were generated by placing the phosphate proton on each of the three oxygen atoms. Hydrogen atoms were added, and protonation states were assigned in MOE. The resulting models were used as initial structures for subsequent DFT calculations.

#### Molecular Dynamics Simulation

All-atom molecular dynamics (MD) simulations were performed using AMBER 24^10^. The ff14SB force field was employed for the proteins, and the TIP3P model was used for water molecules along with Joung-Cheatham parameters for monovalent ions^11–13^. For the modified residues, sTyr and pTyr, force-field parameters were generated using the General AMBER Force Field 2 (GAFF2)^14^. Partial atomic charges (RESP charges) were calculated using the antechamber module based on electrostatic potentials evaluated at the HF/6-31G(d) level in Gaussian^15^. The net charges were set to −1 for sTyr (–SO_3_^−^) and the monoanionic form of pTyr (–HPO_3_^−^), and −2 for the dianionic form of pTyr (–PO_3_^2−^). For monoanionic pTyr, the protonation-site isomer corresponding to the lowest Gibbs free energy of the PSG2-Fv complex, as determined by the DFT calculations described below, was used in the MD simulations. Each complex system was solvated in an orthorhombic box of water molecules with a 30.0 Å buffer from the solute. The systems were neutralized and adjusted to a salt concentration of approximately 45 mM by adding 20 Na^+^ ions and corresponding Cl^−^ ions. The systems were initially energy-minimized and then equilibrated for 1.0 ns under NVT and NPT conditions at 300 K and 1 atm, during which positional restraints on the solute heavy atoms were gradually released. Production MD simulations were carried out for 10 ns under NPT conditions. The temperature was maintained at 300 K using a Langevin thermostat, and the pressure was kept at 1 atm using isotropic pressure scaling. The equations of motion were integrated with a time step of 2 fs, with all bonds involving hydrogen atoms constrained using the SHAKE algorithm. The long-range Coulomb energy was evaluated using the particle mesh Ewald (PME) method with a non-bonded cutoff distance of 9.0 Å. To evaluate the local hydration networks, 200 structural frames were extracted from the production trajectories. For each frame, a hydration subset was defined to include the modified ligand, hydrogen-bonding water molecules identified using a distance cutoff of 3.8 Å and an angle cutoff of 75.0°, and any proximal ions within a 5.0 Å radius. For the dianionic pTyr system (−PO_3_^2−^), the Na^+^ ion closest to the phosphate group in each selected frame was retained in the hydrated configuration used for subsequent DFT calculations.

#### Density Functional Theory (DFT) Calculations

All quantum-chemical calculations were performed with Gaussian 16 using the B3LYP functional with the Grimme’s D3 dispersion correction and the 6-31G** basis set for geometry optimizations and frequency analyses^16^. The initial structure of the PSG2-Fv–sulfotyrosine complex (-SO_3_^−^) was taken from the crystal structure. PSG2-Fv–phosphotyrosine complexes (-HPO_3_^−^, -PO_3_^2−^) were built from the molecular models described above. For monoanionic phosphotyrosine, the three protonation-site isomers were independently optimized within the PSG2-Fv binding-site model, and their Gibbs free energies were compared. The lowest-energy protonation pattern was selected for the MD simulations and subsequent hydrated ligand cluster calculations. The optimized isomers and their relative Gibbs free energies, with the lowest-energy isomer set to 0 kcal mol^−1^, are shown in Figure S11. For the dianionic phosphotyrosine state, one Na^+^ counterion was manually placed near the phosphate group in both the PSG2-Fv complex and isolated-ligand models and was subsequently optimized by DFT. For gas-phase binding free energy calculations, PSG2-Fv binding-site models were generated from the ligand-bound complex structures by extracting the ligand and surrounding residues within 8 Å of the ligand. The resulting binding-site models contained approximately 200 atoms and were used as representative models for DFT calculations. To capture first-sphere hydration effects, hydrated ligand clusters were constructed from MD-derived configurations. From 200 MD-derived frames, 100 representative hydrated configurations were selected for each ligand state. In each configuration, the ligand and its 11 nearest water molecules were extracted to generate ligand·(H_2_O)11 clusters. For dianionic phosphotyrosine, the Na^+^ ion retained from the corresponding MD-derived configuration was also included, generating Na^+^·ligand·(H_2_O)_11_ clusters. This cluster size was chosen to represent a saturated first hydration sphere, consistent with coordination numbers of approximately 11 reported for sulfate and phosphate ions in ab initio MD studies^17,18^. All hydrated ligand clusters were subjected to DFT geometry optimization and frequency analysis. Protein–ligand binding-site models and hydrated ligand clusters were optimized without symmetry constraints. Frequency analyses were performed to obtain thermal corrections to Gibbs free energy at 298.15 K. For optimized structures, the absence of imaginary frequencies confirmed that the geometries corresponded to minima on the potential-energy surface. Thermochemical quantities were evaluated at 298.15 K and 1 atm. Here, complex denotes the optimized PSG2-Fv binding-site model bound to the ligand, derived either from the crystal structure of the PSG2-Fv–sulfotyrosine complex or from the phosphotyrosine-bound molecular models described above. For the dianionic phosphotyrosine state, the complex model also contained one Na^+^ counterion. Fv denotes the corresponding PSG2-Fv binding-site model with the ligand removed and optimized at the same level of theory. Ligand denotes the isolated sulfotyrosine (- SO3-) or phosphotyrosine (-HPO_3_^−^, -PO_3_^2−^). For the dianionic phosphotyrosine state, the ligand model also contained one Na^+^ counterion. Ligand+waters denotes the DFT-optimized cluster composed of the ligand and its 11 nearest water molecules, initially extracted from MD-derived configurations. For the dianionic phosphotyrosine state, this cluster additionally contained the Na^+^ ion retained from the corresponding MD frame. Waters denotes the 11-water cluster obtained from the optimized ligand+waters geometry after removal of the ligand and, for the dianionic phosphotyrosine state, the Na^+^ counterion. The Gibbs free energy of the water cluster was evaluated by frequency analysis at the same level of theory using the inherited geometry, without further geometry optimization. Binding free energies were computed as follows.

*in vacuo*

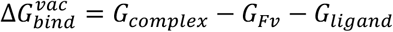

*in aqueous solution*

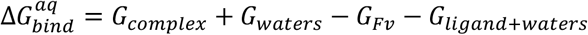

Desolvation free energy for each microstate was computed as follows.

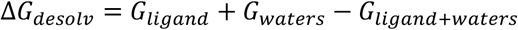

All free energies (*G*) include electronic energy with the thermal correction at 298.15 K and are reported in kcal·mol^−1^. For each ligand, we analyzed an ensemble of 100 distinct hydrated configurations comprising the ligand and its 11 nearest water molecules, summarizing the distribution by the mean and confidence intervals.

#### Differential Scanning Fluorometry (DSF)

DSF data were measured using a CFX Connect Real-Time PCR system (Bio-Rad, Hercules, CA, USA). All scans were obtained at PSG2-Fv concentration of 5 μM in Tris-HCl buffer (pH 7.5) containing 200 mM NaCl and SYPRO-Orange (Sigma, St Louis, MO, USA). Scans were measured from 25 °C to 95 °C at a scanning rate of 1.0 °C/min.

#### Isothermal titration calorimetry (ITC)

The thermodynamic parameters of the interactions between PSG2-Fv and sulfotyrosine or phosphotyrosine were measured with a nano iTC (TA Instruments). Stock solution of sulfotyrosine and phosphotyrosine were quantified by using NMR with the internal standards of the quantification NMR grade DSS (Wako Chemical). A 26 µM solution of PSG2-Fv was titrated with 450 µM sulfotyrosine peptide or 1.1 mM phosphotyrosine peptide in 50 mM Tris-HCl buffer (pH 7.5) containing 200 mM NaCl at 25°C. All the ITC data were analysed using Nanoanalyze software (TA Instruments) with a One Set Sites Fitting Model.

#### Statistical analysis

All analyses were performed in R 4.5.1 (R Foundation). Data wrangling/plotting used tidyverse/ggplot2; inference used rstatix and figure annotation used ggpubr. Unless stated otherwise, tests were two-sided with α = 0.05. Results are reported as means with 95% confidence intervals (CI) calculated from n = 100 representative solvated configurations per ligand, which were representatively selected from the molecular dynamics trajectories. Individual observations are shown as jittered points. Outliers (1.5×IQR) were included in inference. For three-group comparisons of binding energies across ligands, within-group normality was checked by the Shapiro–Wilk test and variance homogeneity by Levene’s test (center = median). Given that the variances were heterogeneous according to Levene’s test (*p* = 1.20 × 10^−7^), Welch’s one-way ANOVA was used to evaluate overall differences among groups, followed by Games–Howell post hoc pairwise tests. Exact and adjusted *p* values, along with *F* statistics and degrees of freedom, are reported in the figures and corresponding legends.

## Supplementary Figures

**Figure S1.**
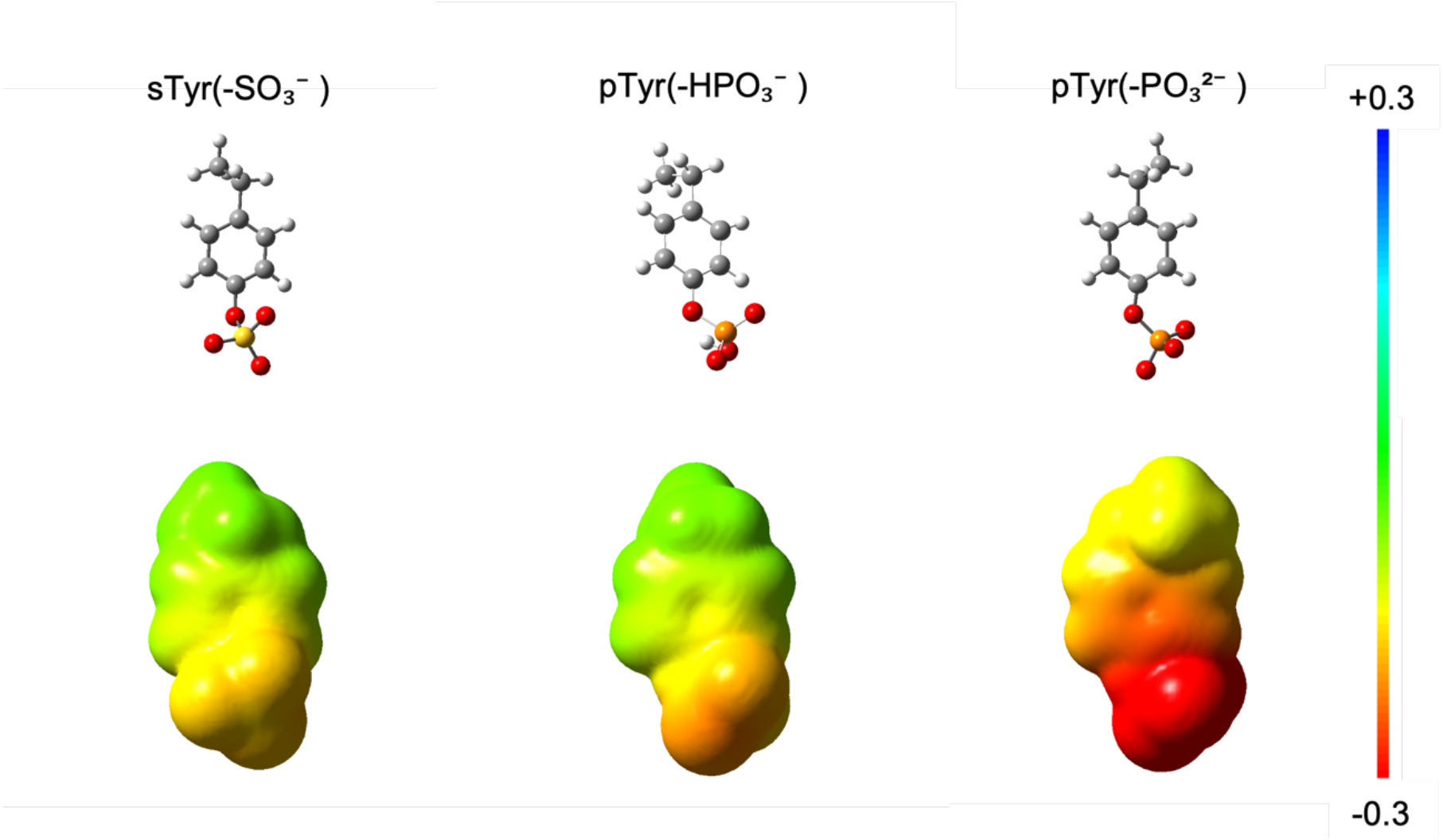
Electrostatic potential (ESP) maps of sTyr and pTyr. Quantum mechanical electrostatic potentials (ESPs) were mapped onto the electron-density isosurface at 0.0004 au, as derived from calculations at the B3LYP-D3/6-31G** level of theory. ESPs are shown on a fixed scale from −0.3 to +0.3 au. Red and blue represent negative and positive electrostatic potential regions, respectively.

**Figure S2.**
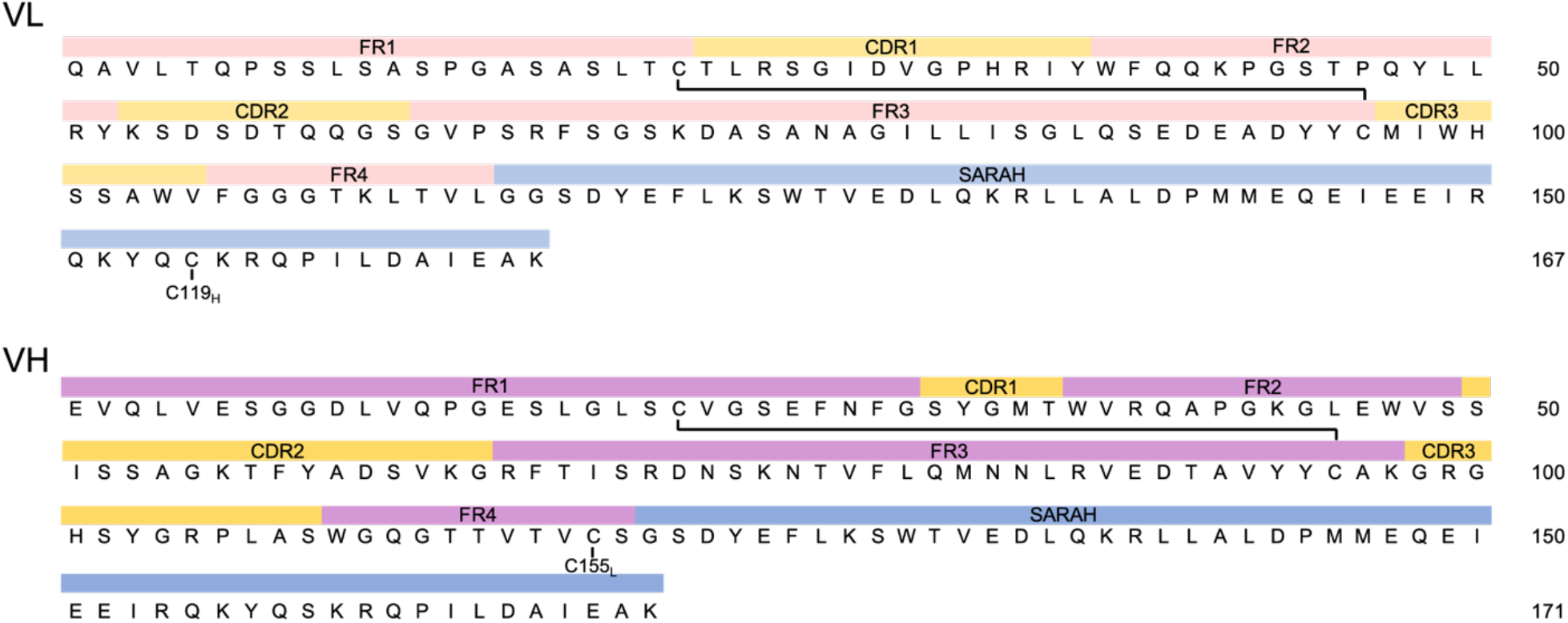
Amino acid sequences of the PSG2 Fv construct. Sequences of the VL (top) and VH (bottom) domains are shown. The framework regions (FR1–4) are shown in pink, the complementarity determining regions (CDR1–3) in yellow, and the SARAH domain in blue. For all regions, light shading is used for the VL domain and dark shading for the VH domain. Intrachain disulfide bonds are indicated by solid lines connecting the respective cysteine residues. The specific cysteine residues introduced for the interchain disulfide bond in the Fv-clasp format are labeled as C119_H_ and C155_L_.

**Figure S3.**
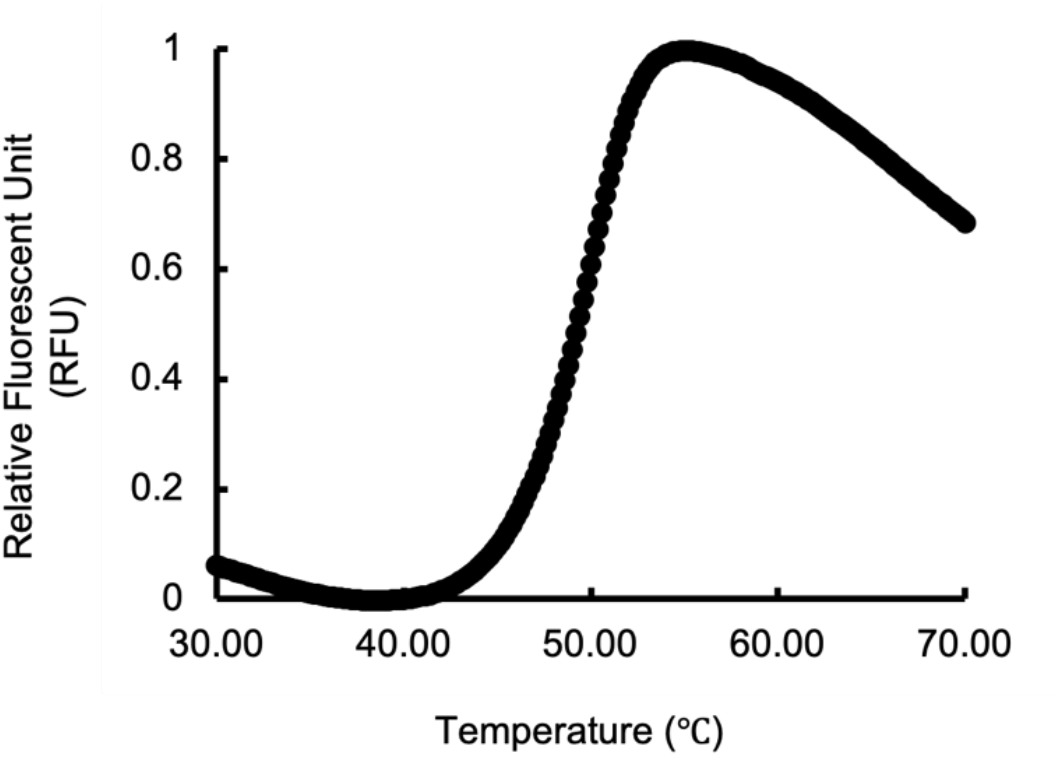
Differential scanning fluorimetry (DSF) analysis of PSG2-Fv in the absence of ligand. The thermal denaturation curve of recombinant PSG2-Fv shows a sigmoidal fluorescence increase with a melting temperature (T_m_) of 49.7℃.

**Figure S4.**
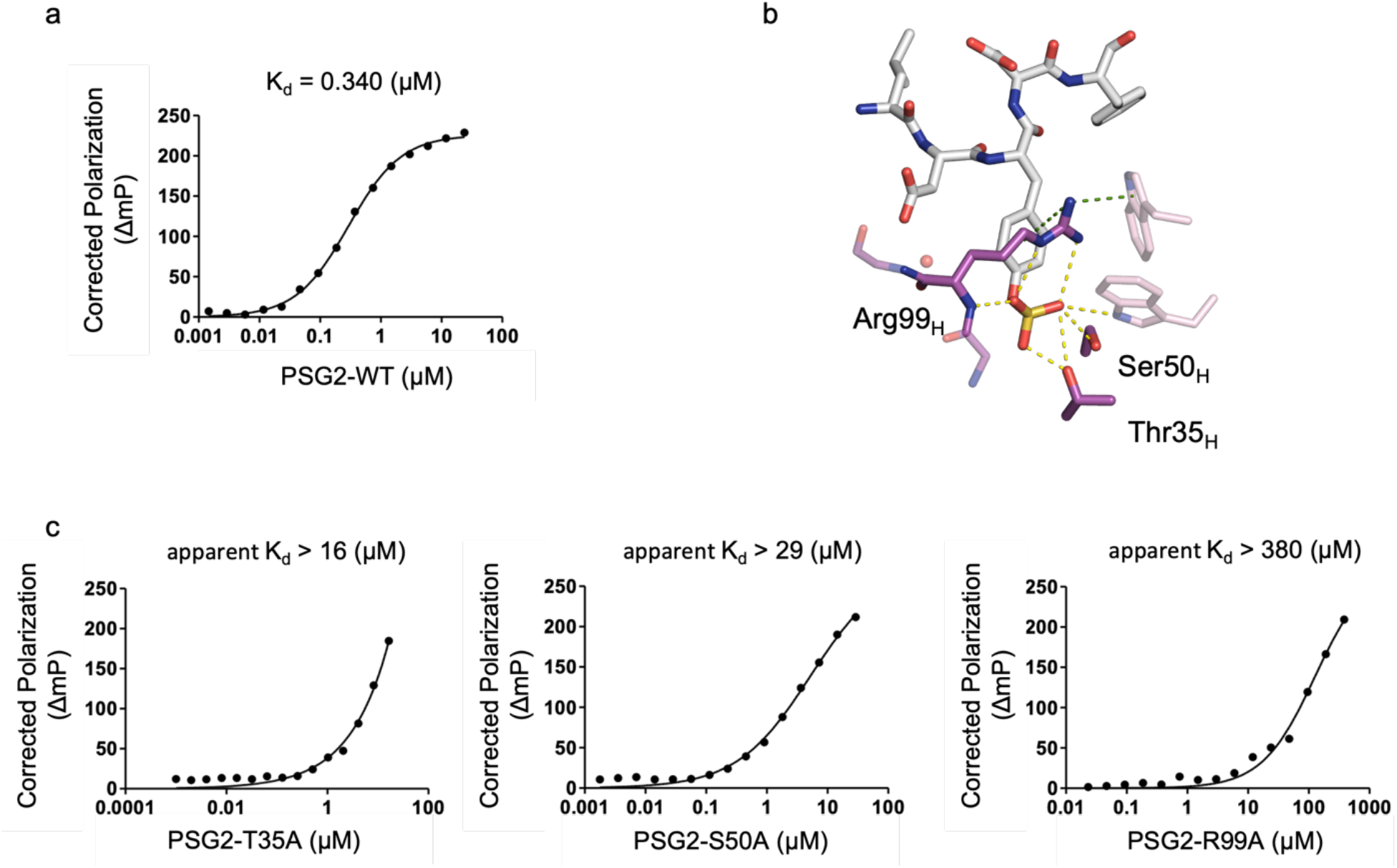
Fluorescence polarization measurements of PSG2-Fv variants. a, Fluorescence polarization measurements of wild-type PSG2-Fv with the fluorescein-labeled sulfotyrosine peptide. b, Structural representation of Thr35, Ser50, and Arg99 in the PSG2-Fv binding pocket. The sulfotyrosine-containing peptide is shown in stick model. The light chain of PSG2-Fv is colored in light pink and the heavy chain of PSG2-Fv colored in magenta. Cation-π interactions are indicated by green dashed lines, and hydrogen bonds are shown by yellow dashed lines. c, Fluorescence polarization measurements of PSG2-Fv mutants (T35A, S50A, and R99A). Measurements were performed once at each PSG2-Fv concentration using different concentration ranges for each mutant. Because saturation was not reached for the mutants within the tested concentration ranges, K_d_ values are reported as lower limits where applicable. The mutants showed reduced fluorescence polarization responses compared with wild-type PSG2-Fv, consistent with weakened binding to the sulfotyrosine-containing peptide.

**Figure S5.**
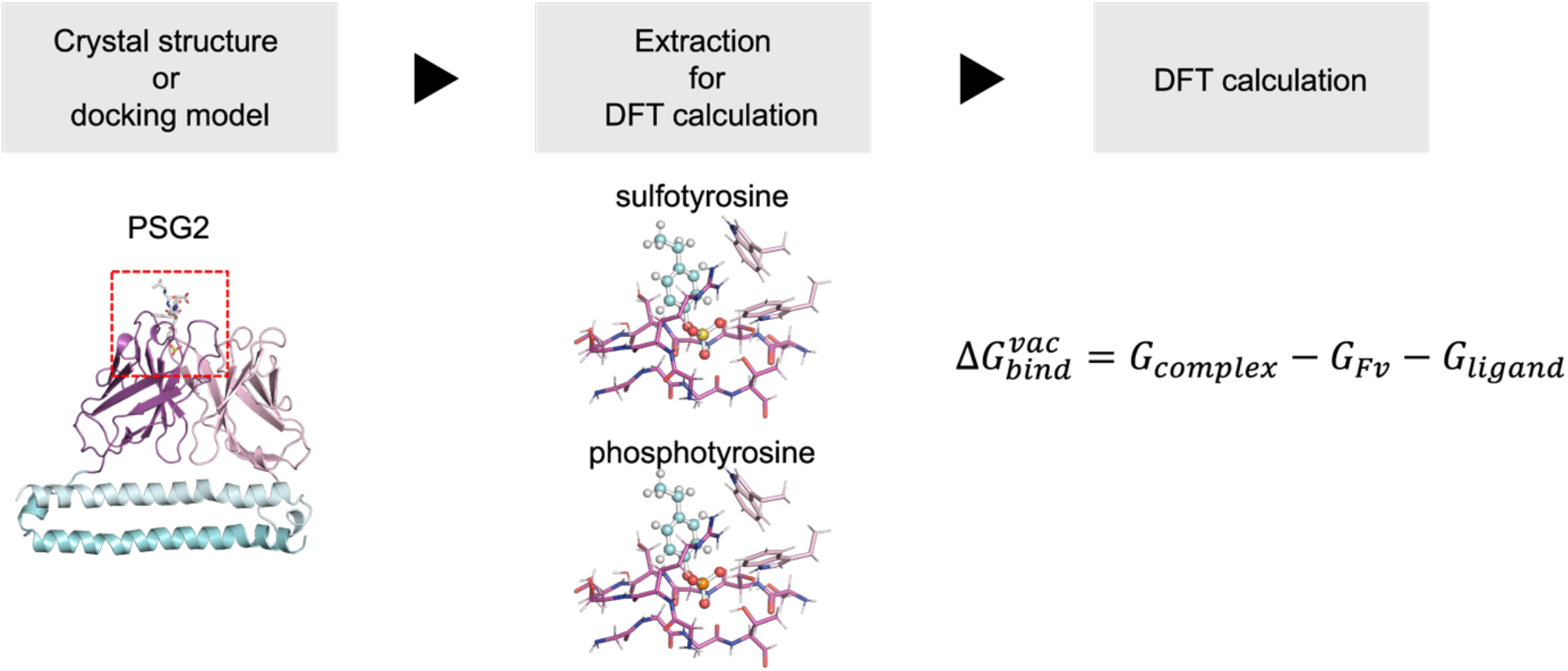
Workflow for *in vacuo* density functional theory calculations of PSG2-Fv bindingsite models. The ligand-bound binding-site models were extracted from the PSG2-Fv–sulfotyrosine crystal structure or phosphotyrosine-bound molecular models and subjected to gas-phase DFT calculations.

**Figure S6.**
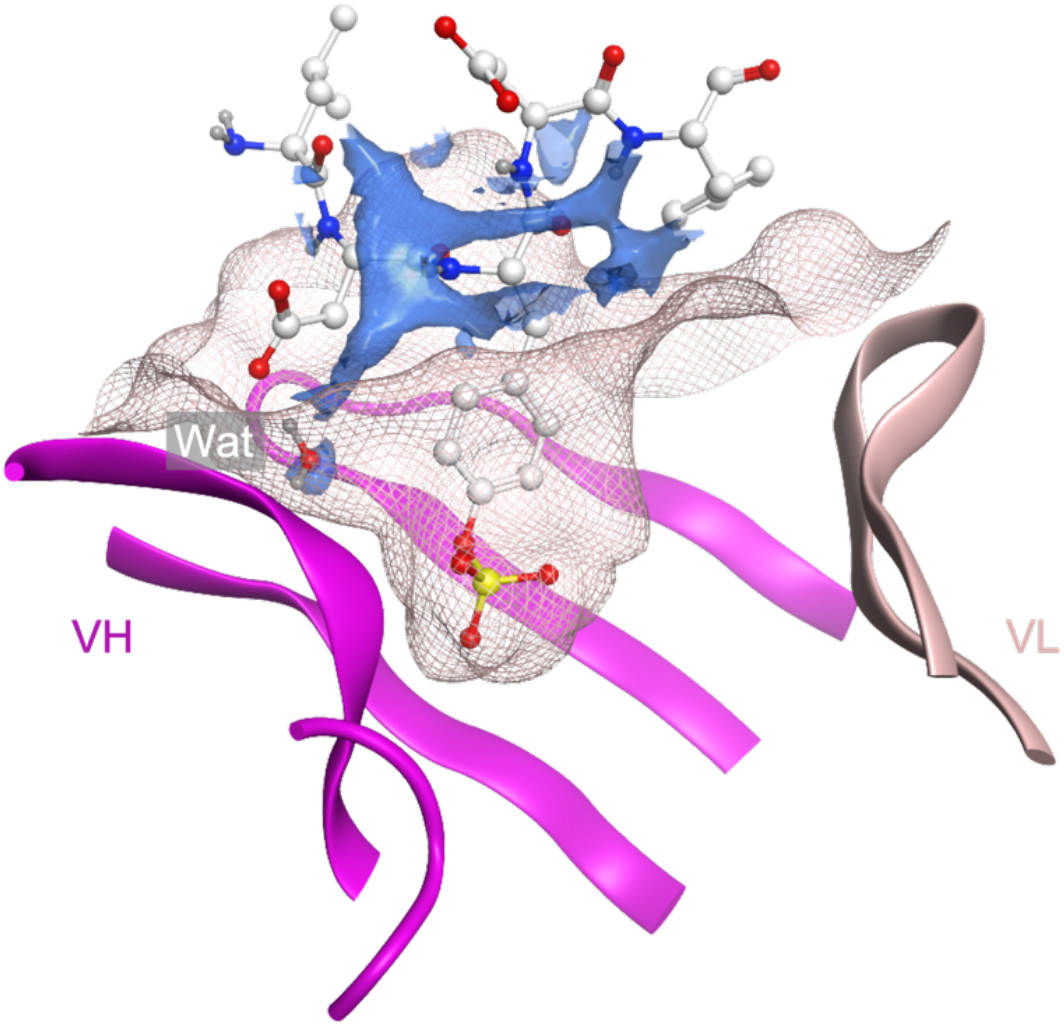
3D-RISM analysis of hydration near the sulfate in the PSG2–sulfotyrosine complex. The blue isosurfaces indicate regions where the calculated water oxygen density is more than four times higher than bulk solvent (relative density > 4.0) within 4 Å of the sulfotyrosine. The PSG2-Fv molecular surface is rendered as a pink mesh, and the sulfotyrosine-containing peptide is shown as sticks. A single water molecule observed in the crystal structure is depicted as a stick, but it is not located near the sulfate. Based on this density threshold, no high-density water site was predicted near the sulfate group.

**Figure S7.**
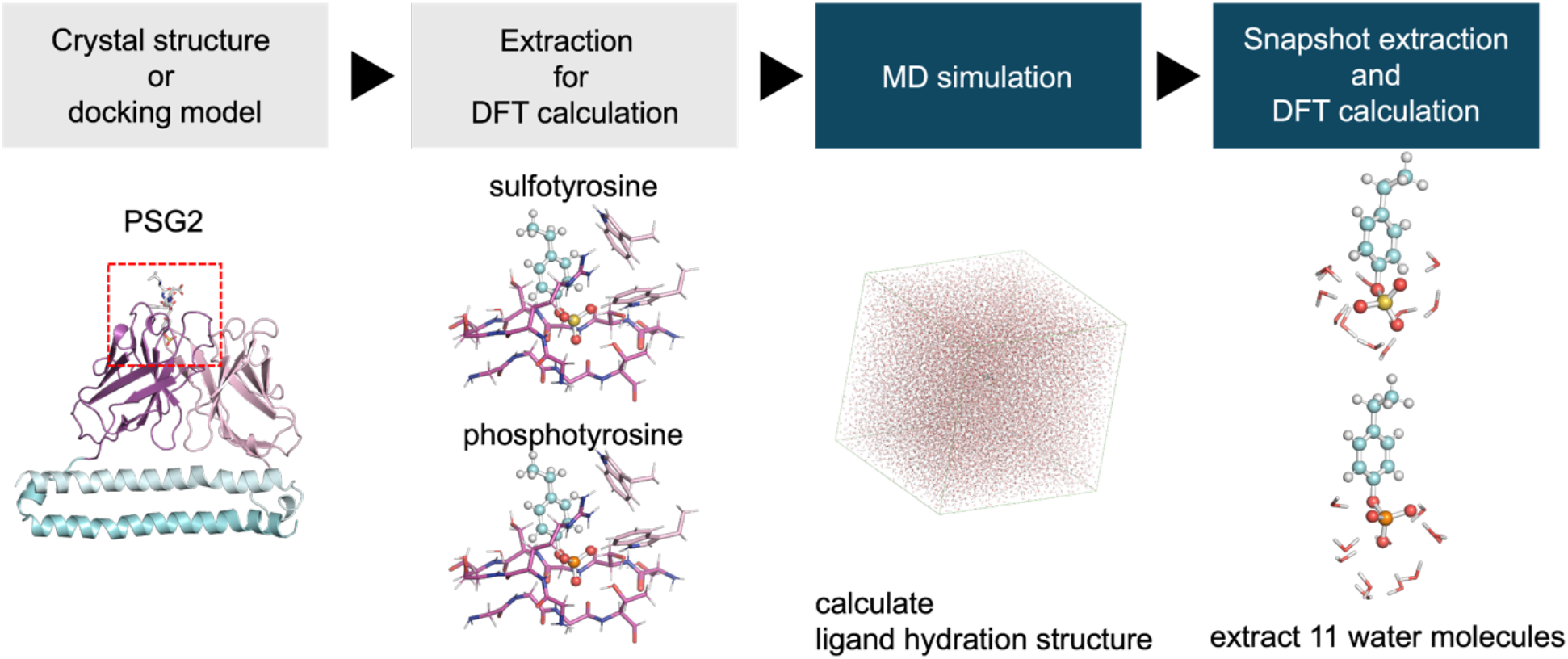
Workflow for constructing hydrated ligand models for DFT calculations. Hydrated ligand clusters were generated from MD-derived configurations by extracting each ligand and its 11 nearest water molecules, and the resulting ligand·(H_2_O)_11_ clusters were used for DFT calculations.

**Figure S8.**
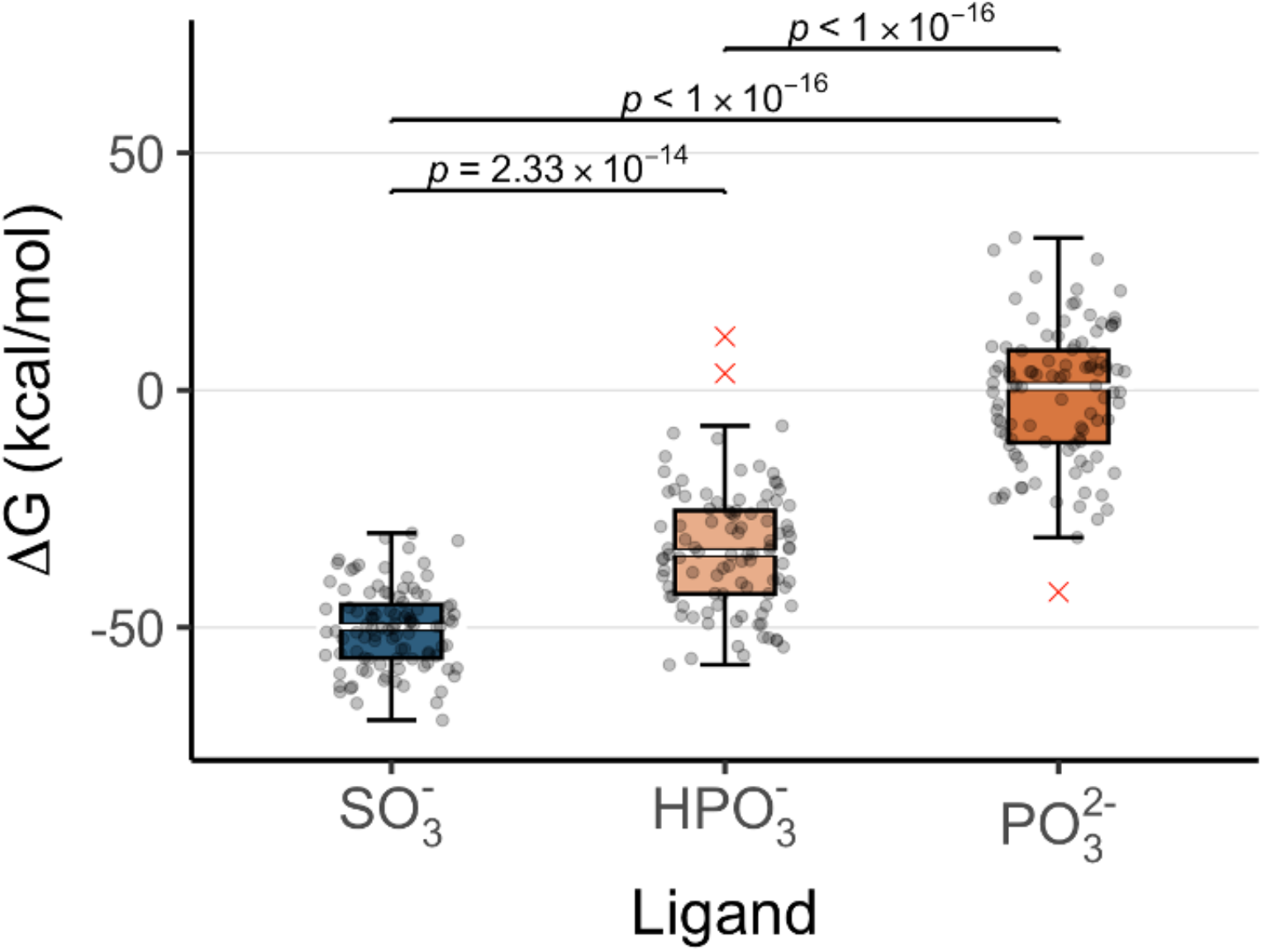
Box plots of binding energies corrected for ligand desolvation. Boxes indicate medians (white line) and IQRs, with whiskers extending to 1.5 × IQR. Individual observations are shown as jittered points (n = 100 ligand·(H_2_O)_11_ cluster models sampled from MD trajectories); outliers (red crosses) were included in all statistical tests. Significant differences among groups were confirmed by Welch’s ANOVA (*F*(2, 186.91) = 428.22, *p* < 0.0001). Adjusted p values from Games– Howell post-hoc tests are annotated in the figure.

**Figure S9.**
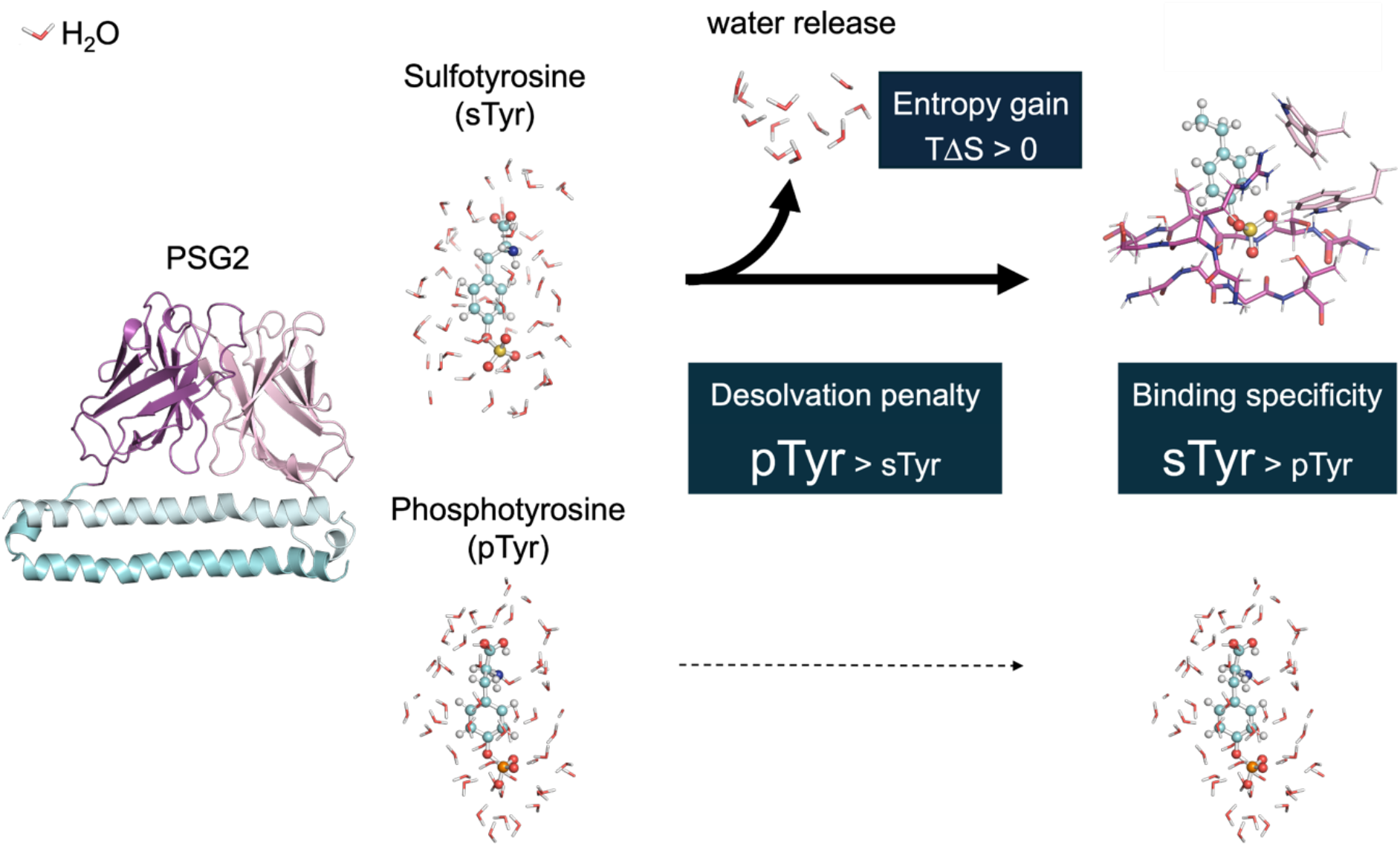
Schematic representation of the PSG2 recognition mechanism for sulfotyrosine (sTyr) over phosphotyrosine (pTyr). The diagram summarizes the structural and energetic factors contributing to PSG2 selectivity. PSG2-Fv is shown as a ribbon model (heavy chain, magenta; light chain, light pink; coiled-coil SARAH domain, light blue). sTyr and pTyr are shown as stick models (carbon, cyan; oxygen, red; sulfur, yellow; phosphorus, orange; nitrogen, blue), together with representative first-shell hydration models containing 11 explicit water molecules. PSG2 preferentially binds sTyr because its smaller desolvation penalty offsets the more favorable direct interactions calculated for pTyr. The favorable entropic contribution observed for sTyr binding may partly reflect the release of water from the ligand and binding pocket.

**Figure S10.**
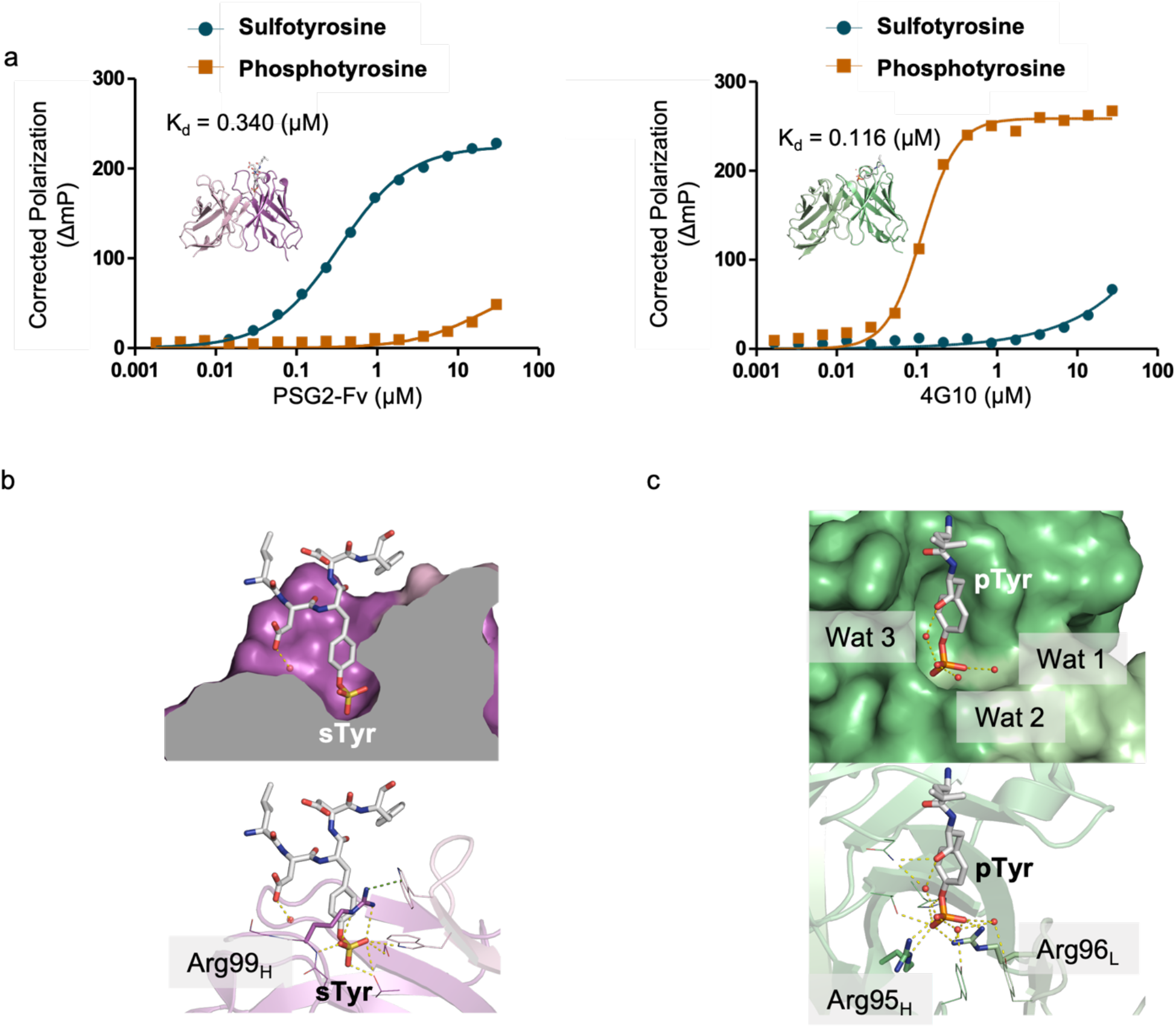
Structural comparison of sulfotyrosine and phosphotyrosine recognition. a, Fluorescence polarization binding assays comparing the ligand preferences of PSG2-Fv and the antiphosphotyrosine antibody 4G10. PSG2-Fv data from Fig. 1b are replotted here for direct comparison with 4G10 and are shown as means ± SEM from triplicate measurements. 4G10 measurements were performed once at each antibody concentration; the K_d_ value shown for 4G10 represents an apparent K_d_ estimated from a single concentration series. PSG2-Fv preferentially bound the sulfotyrosine-containing peptide, whereas 4G10 preferentially bound the phosphotyrosine-containing peptide under the assay conditions used. b, PSG2-Fv bound to an sTyr peptide, showing a deeper, water-poor binding pocket with no modeled water molecules near the sulfate group. Cation–π interactions are indicated by green dashed lines, and hydrogen bonds are shown by yellow dashed lines. c, Anti-pTyr antibody 4G10 (PDB: 6DF1) bound to a pTyr peptide. The phosphate group is coordinated by multiple hydrogen bonds and salt bridges from conserved cationic residues, while three water molecules remain in the binding site. Hydrogen bonds are shown by yellow dashed lines.

**Supplementary Fig. 11.**
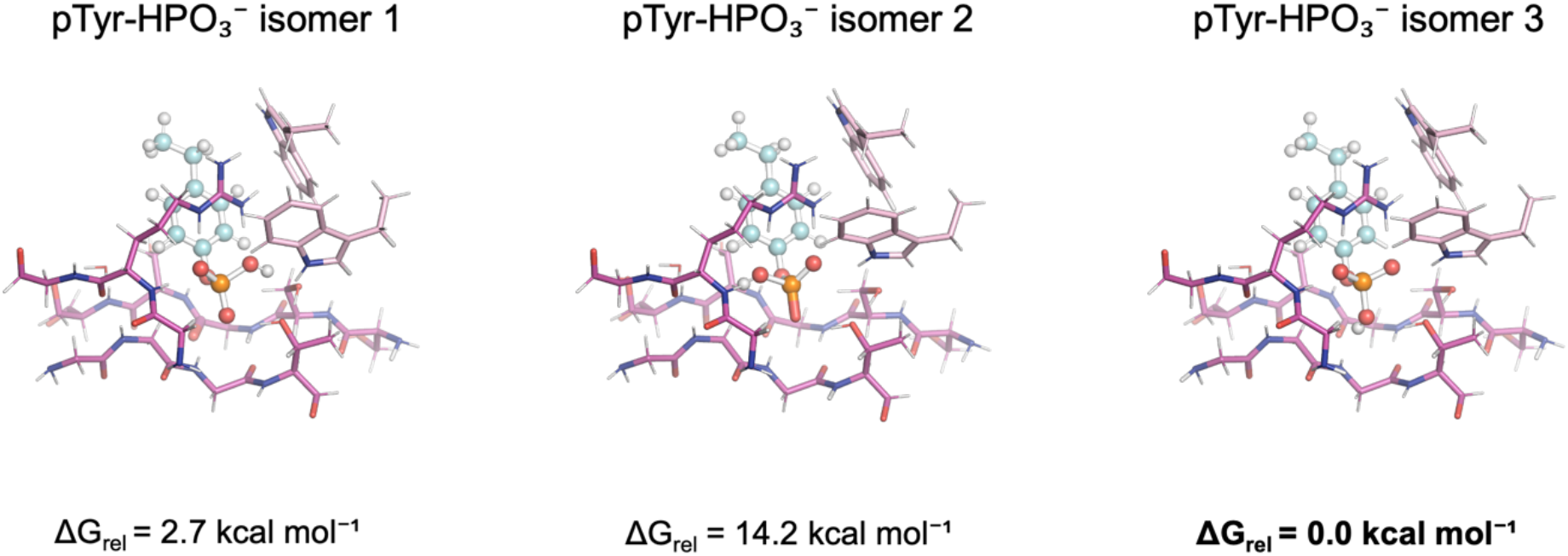
DFT-optimized structures of protonation-site isomers of monoanionic phosphotyrosine in the PSG2-Fv binding-site model. Three protonation-site isomers of monoanionic phosphotyrosine (−HPO_3_^−^), differing in the position of the phosphate proton, were independently optimized using DFT at the B3LYP-D3/6-31G** level. Relative Gibbs free energies (ΔG_rel_) are shown in kcal mol^−1^, with the lowest-energy isomer set to 0.0 kcal mol^−1^. The lowest-energy protonation pattern was used for the subsequent MD simulations and hydrated ligand-cluster calculations.

## Supplementary Tables

**Table S1.** Amino acid sequences of PSG2-Fv expression constructs.

|  |
| --- |
| **PSG2-Fv_VL** |
| MGHHHHHHGGQAVLTQPSSLSASPGASASLTCTLRSGIDVGPHRIYWFQQKPGSTPQYLLRYKS<br>DSDTQQGSGVPSRFSGSKDASANAGILLISGLQSEDEADYYCMIWHSSAWVFGGGTKLTVLGGS<br>DYEFLKSWTVEDLQKRLLALDPMMEQEIEEIRQKYQCKRQPILDAIEAK |
| **PSG2-Fv_VH** |
| MSKIKGHHHHHHGGEVQLVESGGDLVQPGESLGLSCVGSEFNFGSYGMTWVRQAPGKGLEWV<br>SSISSAGKTFYADSVKGRFTISRDN SKNTVFLQMNNLRVEDTAVYYCAKGRGHSYGRPLASWGQ<br>GTTVTVCSGSDYEFLKSWTVEDLQKRLLALDPMMEQEIEEIRQKYQSKRQPILDAIEAK |

**Table S2.**
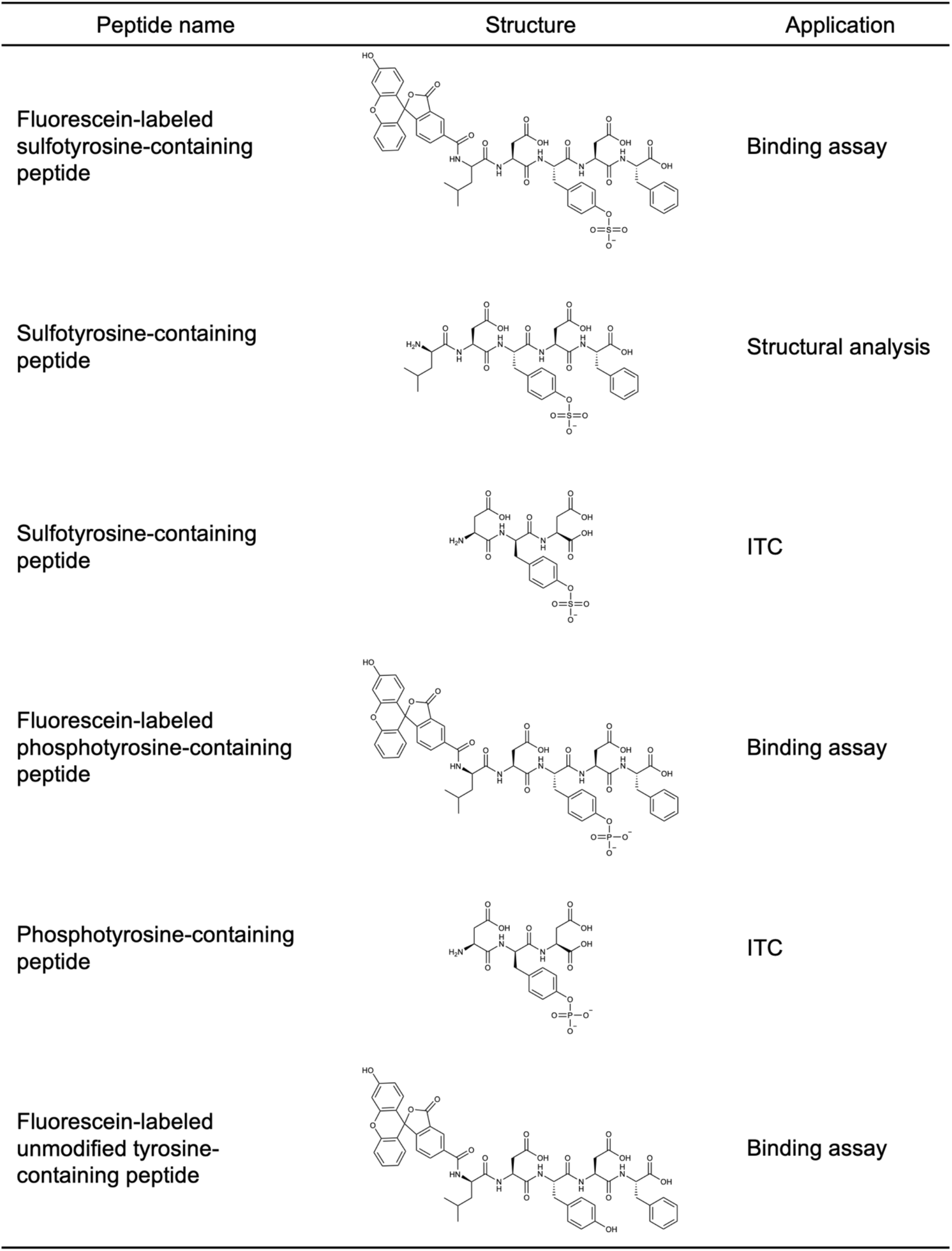
Synthesized peptides and their experimental applications.

| Peptide name | Structure | Application |
| --- | --- | --- |
| Fluorescein-labeled sulfotyrosine-containing peptide |  | Binding assay |
| Sulfotyrosine-containing peptide |  | Structural analysis |
| Sulfotyrosine-containing peptide |  | ITC |
| Fluorescein-labeled phosphotyrosine-containing peptide |  | Binding assay |
| Phosphotyrosine-containing peptide |  | ITC |
| Fluorescein-labeled unmodified tyrosine-containing peptide |  | Binding assay |

**Table S3.**
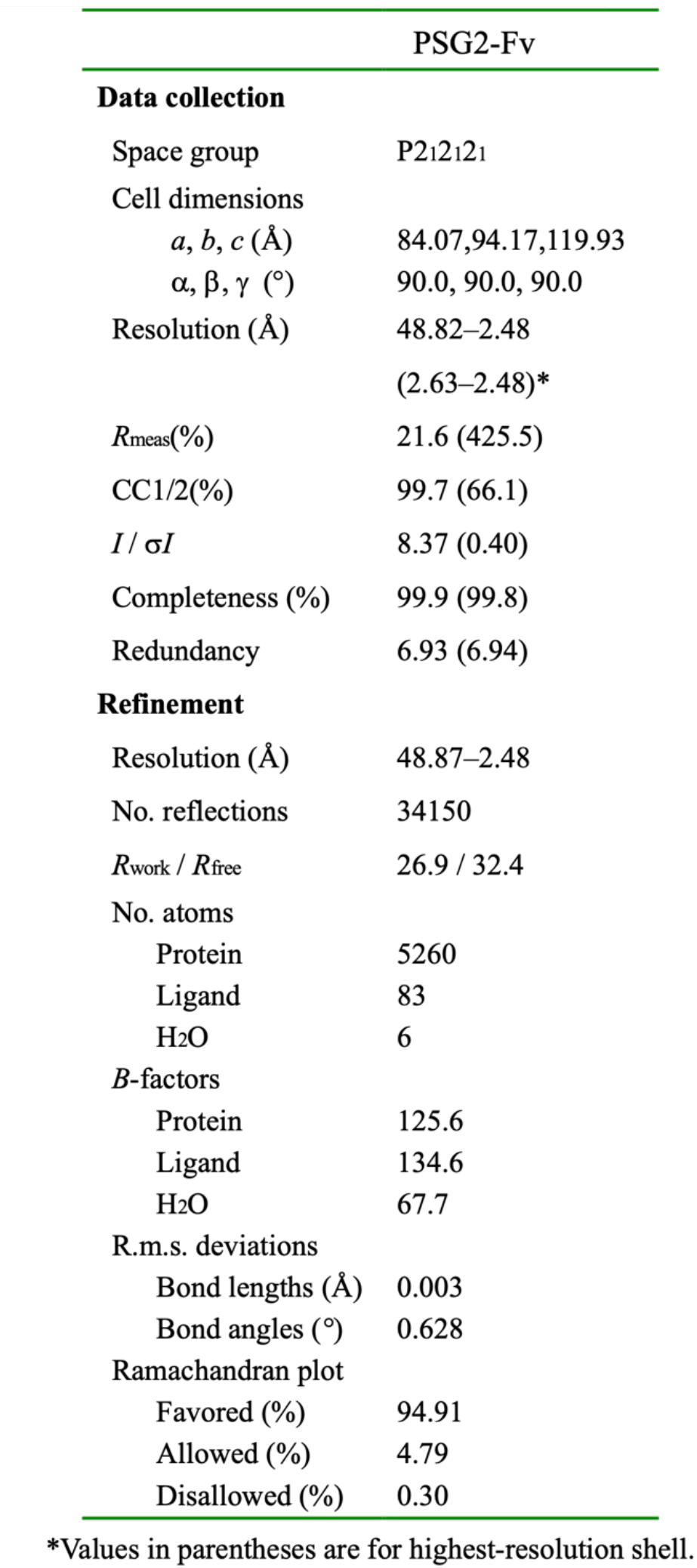
Crystallographic data collection and refinement statistics.

## Notes

### Competing Interest Statement

The authors have declared no competing interest.

